# Motor Evoked Potentials in Preclinical Models of Spinal Cord Injury and Stroke in Rodents: A Systematic Review

**DOI:** 10.64898/2026.10.09.757956

**Authors:** Rubab Shahid, Dylan F. Cooke, Gustavo Balbinot

## Abstract

Motor Evoked Potentials (MEPs) serve as an invaluable tool for assessing corticospinal tract (CST) integrity and motor pathway function in preclinical models of spinal cord injury (SCI) and stroke. Despite their diagnostic and prognostic utility, substantial methodological variability persists across studies, complicating interpretation and cross-study comparisons. This systematic review synthesizes MEP acquisition and interpretation protocols from 83 rodent SCI and stroke studies published since 2015, aiming to outline common practices, identify sources of variability, and highlight methodological gaps. Our analysis reveals a predominance of SCI models and a focus on hindlimb MEPs, employing diverse stimulation protocols including transcranial electrical stimulation (TES), direct cortical stimulation (DCoS), and transcranial magnetic stimulation (TMS). Anesthetic approaches are notably heterogeneous, representing a potentially large source of variability in MEP acquisition. Amplitude and latency, often measured jointly, emerged as the most frequently reported MEP parameters. Critically, MEPs bridge cellular and molecular changes to observable behaviour by reflecting the integrated functionality of neural cell populations and their circuits. However, a lack of standardization in reporting, particularly for secondary electrophysiological features and stimulation intensity, hinders reproducibility. We emphasize the necessity for standardized protocols for stimulation, electrode positioning, and data analysis, particularly the use of comprehensive recruitment curves. Adopting these best practices will enhance the transparency, comparability, and translational potential of MEP methodology in advancing our understanding and treatment strategies for neurological disorders.

**Graphical abstract:** 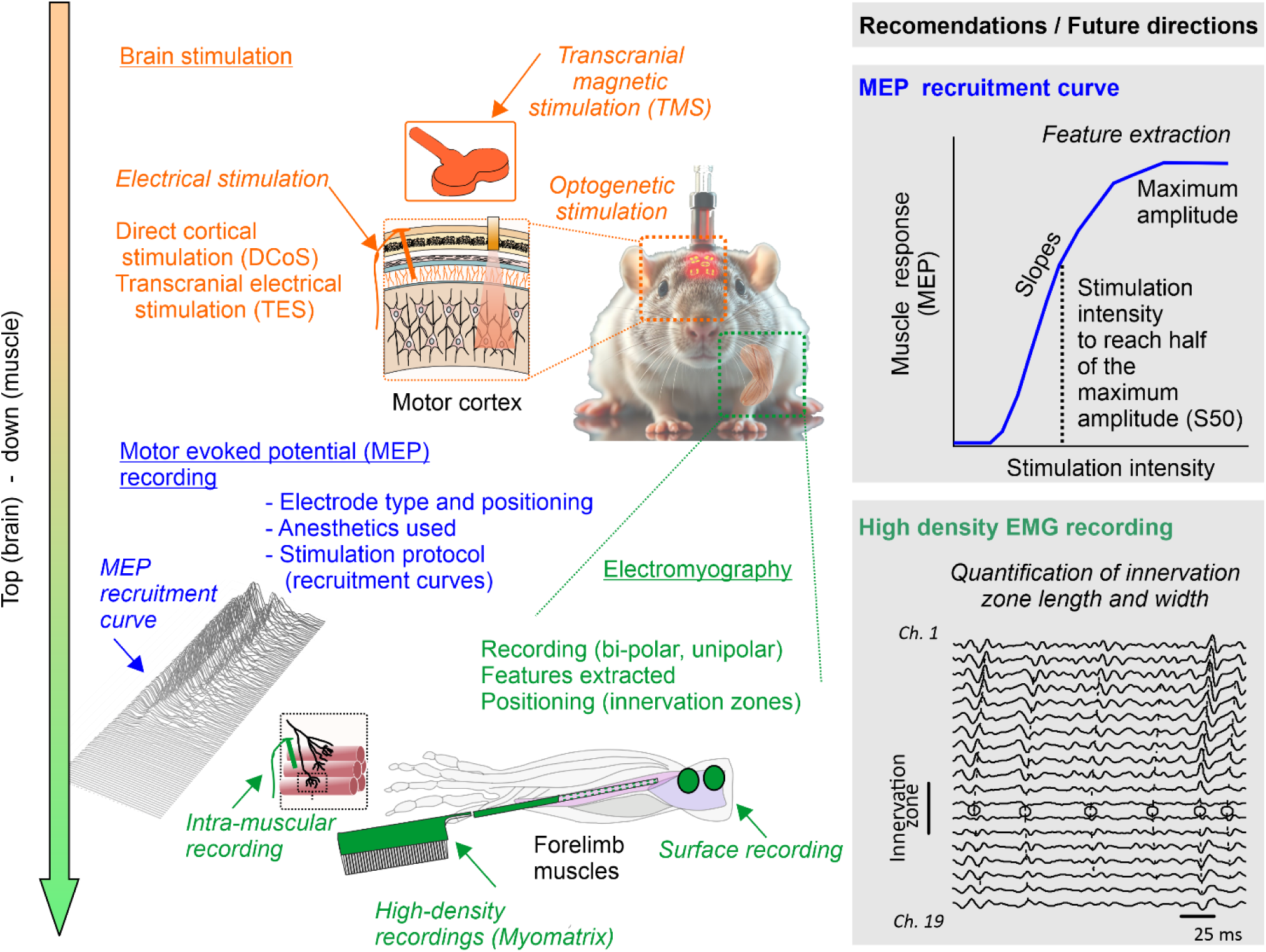

**Motor evoked potentials (MEPs) in rodent models of spinal cord injury (SCI) and stroke.** This review focuses on various brain stimulation techniques (orange) and electromyographic approaches employed to quantify MEPs (green) and factors affecting MEP recording and analysis (blue). Future directions include greater standardization of stimulation, recording-electrode placement, and data-analysis practices; such as the potential use of recruitment curve features and high-density EMG recordings such as Myomatrix. Recruitment curves provide a more comprehensive characterization of evoked responses than single-pulse stimulation. While a single pulse only offers a snapshot of a subject’s motor threshold, recruitment curves yield a dynamic, S-shaped (sigmoidal) function that reflects the net excitability of upper and lower motor pathways across a range of stimulation intensities. This enables better control over sources of variability, such as stimulation and recording electrode placement, anesthetics dose, that can alter amplitude-based features. Because recruitment curves quantify the relationship between stimulus intensity and response rather than relying solely on absolute amplitude, the derived features are less susceptible to these artifacts (e.g., slopes, S50—stimulation intensity to reach half of the maximum amplitude). In addition, high-density recordings make it possible to better characterize the evoked activity of multiple motor units within the same target muscle and to identify the innervation zones of the motor units under study.

*TMS (Transcranial magnetic stimulation); DCoS (Direct cortical stimulation); TES (Transcranial electrical stimulation); MEP (motor evoked potential); Ch. (EMG channel)*.

## 1. Introduction

Loss of motor function is one of the most debilitating outcomes of stroke and spinal cord injury (SCI), severely limiting independence and quality of life. These deficits primarily arise from loss of cortical motor circuits and disrupted signal transmission along the corticospinal tract (CST), a major descending motor pathway with limited regenerative capacity within the central nervous system (CNS) (1–4). Indeed, residual CST integrity is a strong predictor of motor recovery and treatment response (2–4). Motor recovery thus depends on both structural preservation and the ability to transmit signals through injured motor pathways; therefore, measuring signal transmission in descending tracts is a key neurophysiological indicator. Consequently, functional assessment of descending pathway integrity is important in both clinical and preclinical neurotrauma research.

Motor evoked potentials (MEPs) provide one method of assessing motor pathway function—transmission of sensorimotor information from the brain. MEPs represent the electromyographic responses recorded from muscles following cortical stimulation and provide a dynamic measure of corticospinal excitability. MEP presence indicates whether a measurable motor response can be elicited (i.e., descending motor output is present), amplitude reflects the magnitude of this output, and latency shows the time required for the signal to reach the target muscle. Together, these MEP features characterize whether descending motor output is present, how strongly it is expressed, and how efficiently signals are conducted to the target muscle.

In clinical populations, MEP presence and selected quantitative measures, particularly amplitude, have been used to predict motor recovery after stroke and SCI, providing functional information that can complement structural imaging (5,6). These measures can also help identify muscle groups with greater recovery potential during early post-injury stages (7,8) and contribute to predictive models of recovery (1,4).

MEP characteristics depend partly on how the motor system is stimulated and the response recorded; see **Box 1** for details on the mode of stimulation and recording.

#### Box 1. Stimulation and recording of MEPs in rodents

**Table A1** introduces the principal stimulation modalities used to elicit motor evoked potentials (MEPs) in rodent models—transcranial magnetic stimulation (TMS), transcranial electrical stimulation (TES), direct cortical stimulation (DCoS), and optogenetic stimulation—and summarizes how they differ in invasiveness, spatial specificity, typical parameterization, and practical trade-offs. Because stimulation choice, physical setup or “montage” (device/electrode type), and waveform (intensity, pulse width, and train structure) strongly shape MEP threshold, amplitude, latency, and waveform morphology, the table highlights expected impacts on MEP readouts to aid method selection and transparent reporting.

**Table A1.** Stimulation modalities overview

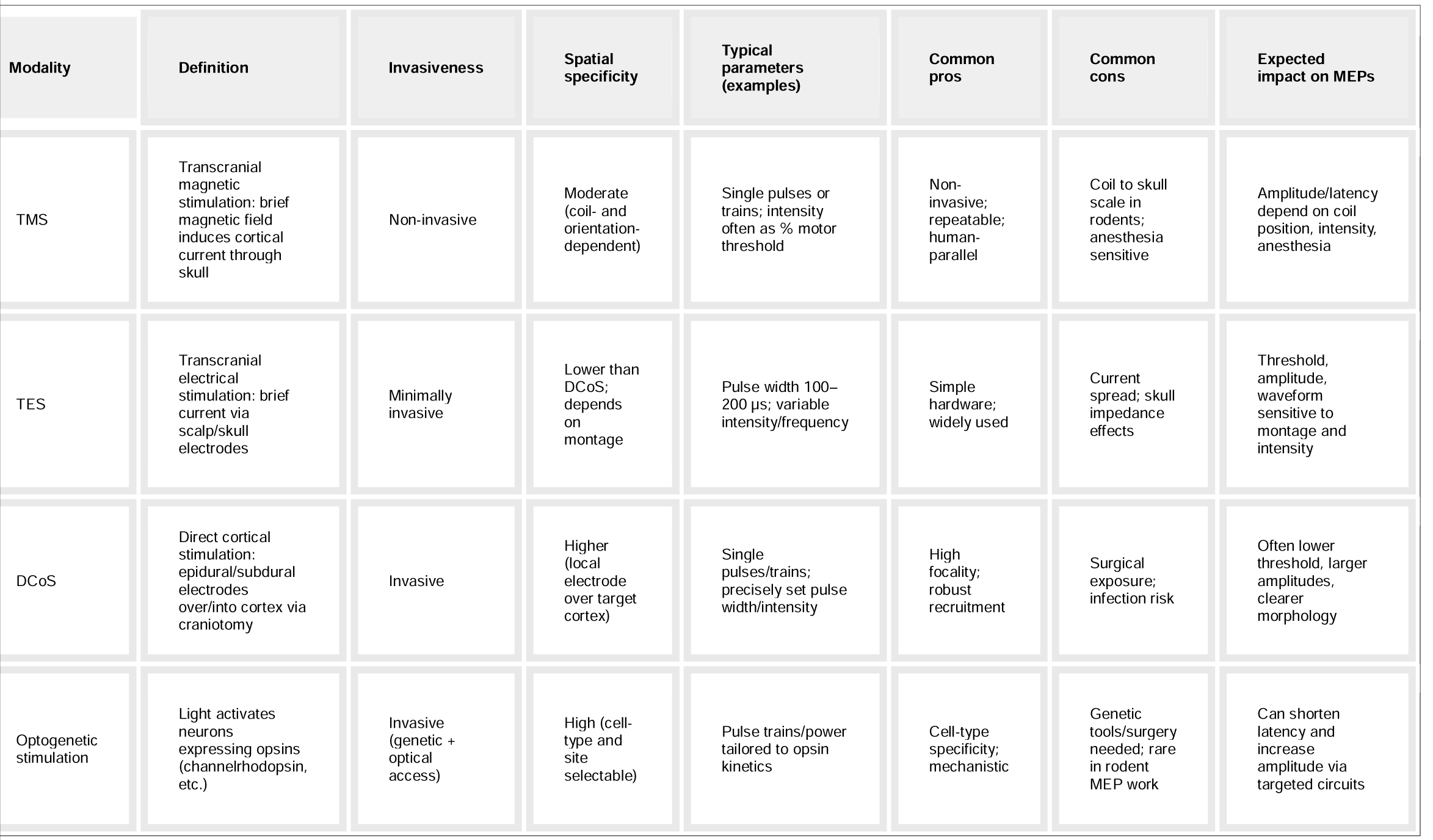

**Table B1** outlines the most commonly used EMG-derived MEP metrics: amplitude, area under the curve (AUC), latency to onset, latency to peak, waveform morphology, and presence/threshold, with concise operational definitions, their physiological interpretation, and practical considerations for reproducible measurement.

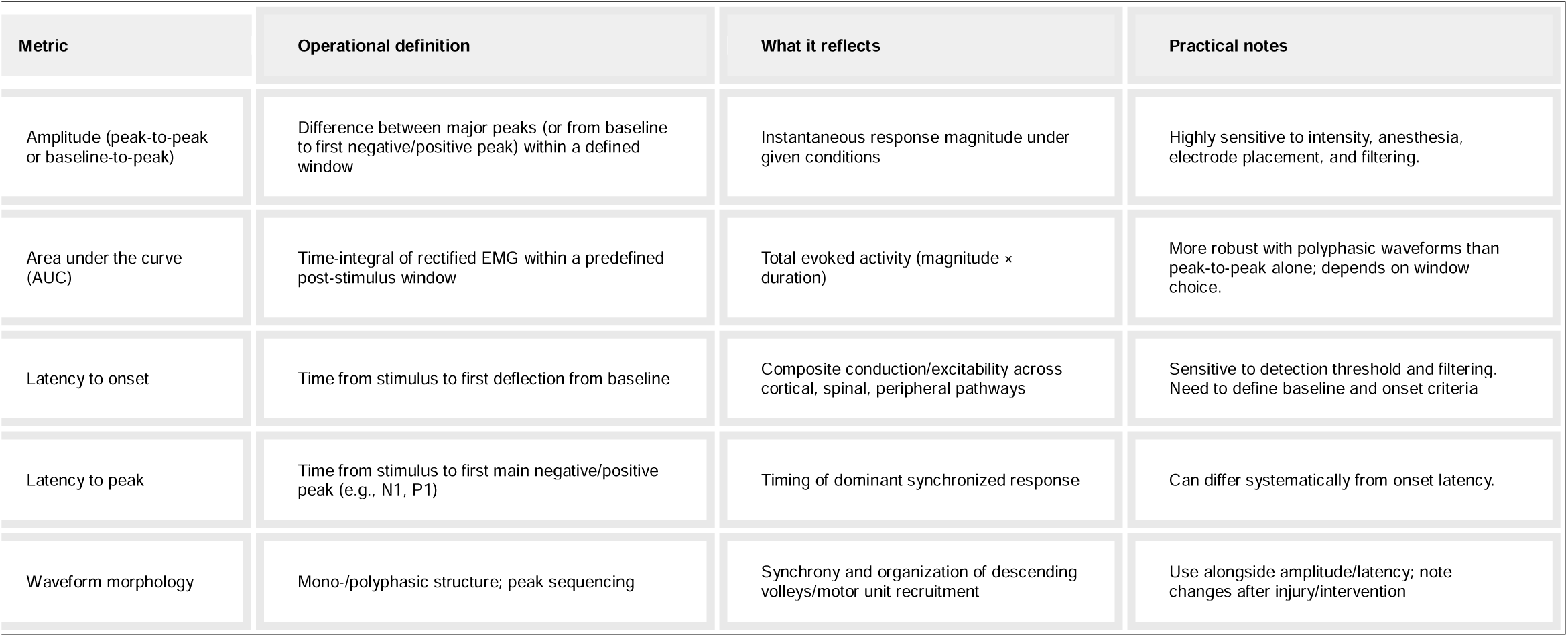

**Figure B1** provides a visual example of how these metrics can be extracted from a representative TMS-evoked MEP waveform.

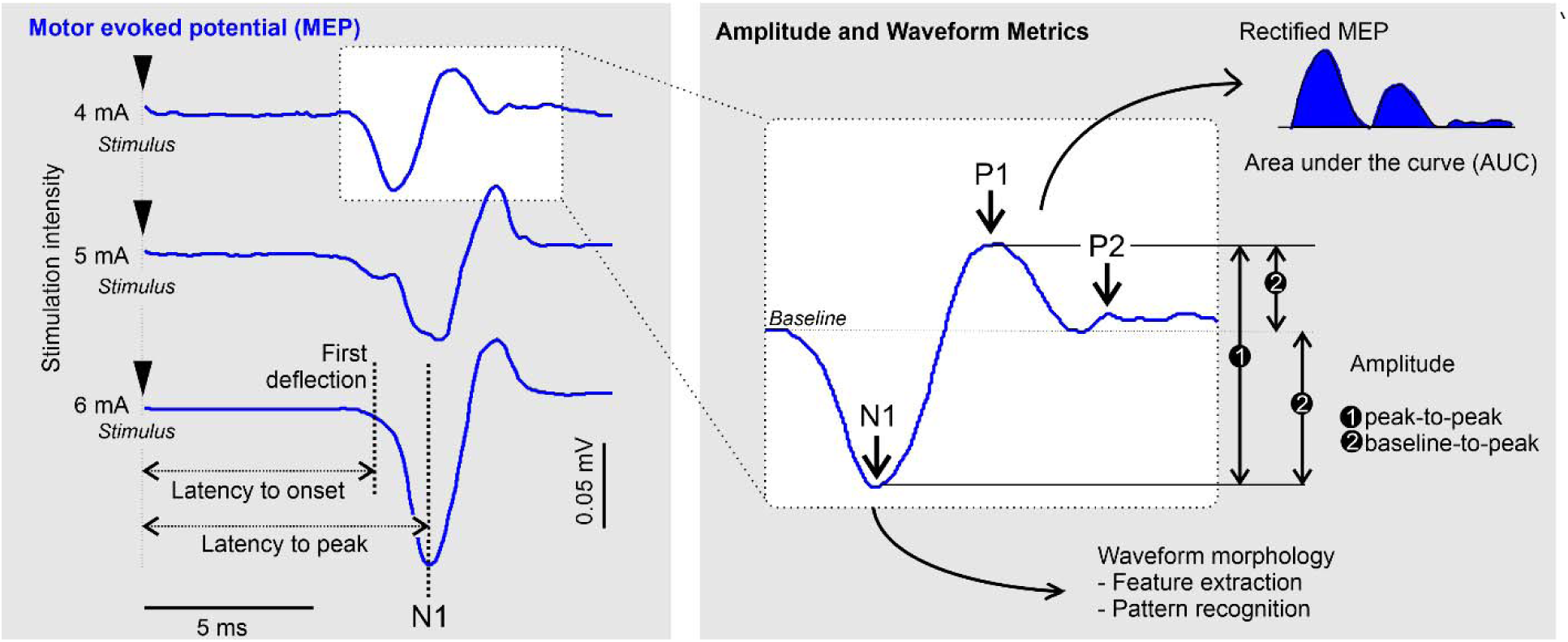

**Figure B1.** Representative TMS-evoked MEP waveforms and extraction of common EMG-derived metrics. Left: Representative MEP traces recorded from rat calf muscle following TMS at increasing stimulation intensities (4–6 mA). The stimulus, first detectable waveform deflection, latency to onset, and latency to the first negative peak (N1) are indicated. Right: Enlarged waveform showing the N1, P1, and P2 phases and examples of baseline-to-peak or peak-to-peak amplitude measurement, rectified area under the curve (AUC), and waveform-morphology analysis through feature extraction and pattern recognition. Waveform traces and N1–P1–P2 nomenclature were adapted from Luft et al. (2001) (9).

Interpretation of MEP outcomes is influenced by the muscle recorded, electrode placement, stimulation mode, anesthesia, and signal-processing methods (10,11). These methodological considerations are particularly important in preclinical stroke and SCI research, where MEPs are used to quantify neurophysiological changes after injury and evaluate interventions intended to promote neural repair or functional recovery. Rodent MEPs complement behavioural, anatomical, and molecular assessments by linking stimulation of the motor system to a measurable downstream muscle response. MEPs provide a crucial link between cellular or pathway-level changes and observable motor behaviour. Behavioural assays (ladder rung, grid walking, cylinder tests, BBB/BMS scales) are valuable but rely on visual scoring and may miss subtle or early recovery (12). Histological and biochemical methods (axon tracing, immunohistochemistry, molecular analyses) reveal cellular and structural changes but do not show whether functional signal transmission is restored. Other electrophysiological tools—somatosensory evoked potentials, compound muscle action potentials, EMG, and nerve conduction studies—assess complementary aspects of sensory and motor pathway function (1,13).

Rodent models provide controlled frameworks for investigating injury mechanisms and testing therapeutic and rehabilitative interventions. In these models, MEPs may detect neurophysiological changes before overt behavioural improvement and help relate cellular or structural findings to functional motor outcomes. Significantly, roundtable meetings identified barriers to translating non-invasive brain stimulation research into practice and recommended improving mechanistic understanding, strengthening preclinical methodological rigor, standardizing outcome measures, and enhancing clinical relevance of animal models (14). MEPs play a crucial role in all of these factors, yet despite their importance, MEP methodologies remain highly variable (10). Differences in stimulation modality, signal processing, anesthesia, feature extraction, and reporting conventions can all influence the resultant MEP signal. These sources of variability complicate interpretation and cross-study comparison, making it difficult to determine whether differences in MEP outcomes reflect true biological effects, methodological differences, or both. It is likely that inconsistent reporting also poses a barrier for new researchers seeking to select MEP methodologies most appropriate for their experimental aims. Reliability studies are important (15), but a systematic review of rodent MEP research would clarify the field’s current status and could foster further consensus and roundtable meetings.

Previous reviews on MEPs have largely focused on assessment of human CST with transcranial magnetic stimulation (TMS), including best practices for obtaining reliable MEP measures (16). The most recent narrative review of rodent MEP techniques was published in 2012 and concentrated specifically on TMS (17), while reviews on rodent models of stroke or SCI have focused on stimulation methods rather than MEP acquisition, quantification, and interpretation (18). An updated systematic review of current MEP practices in rodent models of stroke and SCI is therefore needed.

This systematic review summarizes MEP acquisition and analysis practices reported in rodent models of stroke and SCI over the past decade. Specifically, we examine study characteristics, intervention categories, stimulation and recording parameters, anesthesia practices, reported MEP outcome measures, and approaches to MEP quantification across different experimental applications. By identifying common practices, reporting gaps and sources of methodological variability, the review aims to support protocol selection and greater standardization in preclinical MEP research. Improving the consistency and interpretability of MEP methodology will strengthen reproducibility and enhance the translational value of electrophysiological measures of motor system recovery.

## 2. Methods (445 words)

### 2.1 Information Sources and Search Strategy

A systematic search was conducted in MEDLINE (via PubMed) to identify studies investigating MEPs in rodent models of SCI or stroke. The search covered publications from January 1, 2015 to the date of the final search—December 16, 2024 (see **Supplementary Materials** for full strategy). The 2015 cutoff was selected to ensure inclusion of contemporary methodological practices in MEP acquisition and interpretation. No restrictions were placed on publication status. Only studies published in English were included. Reference lists of all eligible studies were screened manually to identify additional publications.

### 2.2 Eligibility Criteria

Studies were included if they met the following criteria. Inclusion criteria: Rodent models (rats or mice; any strain, sex, or age); Experimental stroke or SCI model; Healthy or sham controls (either as separate groups or via pre–post comparison); Reported MEPs and described acquisition methodology. Exclusion criteria: Non-rodent; ex vivo; in vitro; human studies; computational models; No pre/post comparison when relevant; Articles published before 2015; Non-English language publications; Reviews, perspectives; conference abstracts; theses; or non-primary sources. The review protocol was prospectively registered in PROSPERO (Registration ID: CRD420250631001).

### 2.3 Study Selection

Search results were imported into a spreadsheet for screening. Two reviewers (GB and RS) independently screened all titles and abstracts (Stage 1), followed by full-text evaluation of potentially eligible studies (Stage 2). Any disagreements were resolved through discussion and consensus; unresolved conflicts were adjudicated by GB.

### 2.4 Data Collection Process

Data were extracted by RS under the supervision of GB using a standardized extraction form developed for this review. Extraction was performed independently and cross-checked for accuracy.

### 2.5 Data Items

The following variables were extracted from each study:

#### 2.5.1 General Study Characteristics

Publication year, study design, sample size. Species, strain, sex, age. Lesion model (stroke or SCI type, severity, location). Study aim.

#### 2.5.2 Experimental Procedures

Post-lesion interventions (e.g., stem cell therapy, biomaterials, neuromodulation, pharmacological treatments). Target muscles recorded. Stimulation modality (electrical, magnetic, or other). Stimulation parameters (intensity, pulse duration, frequency, electrode type). Electromyography (EMG) recording setup. Anesthetic(s) used during MEP acquisition.

#### 2.5.3 MEP Measurement and Signal Processing

MEP properties analyzed (amplitude, latency, area under the curve, signal presence, waveform morphology). Electrode placement (stimulation and recording sites). Signal filtering, artifact removal, averaging, and threshold criteria. Hardware and software used.

#### 2.5.4 Outcomes

MEP outcomes (quantitative and qualitative). Behavioural or functional motor outcomes when reported.

### 2.6 Synthesis Methods

Due to substantial methodological variation, findings were synthesized descriptively. Extracted variables were grouped into thematic domains: (1) study characteristics, (2) intervention categories, (3) MEP quantification practices, (4) methodological reporting quality, and (5) anesthesia practices. No meta-analysis was performed. The extracted variables formed the basis for the methodological synthesis reported in the Results section. We used the following nomenclature: NA for information not available (*e.g*., there was no intervention) and NDP for information available but with no details provided.

## 3. Results (1174 words)

### 3.1. Analysis

**Figure 1** shows the flowchart of the literature search strategy and study screening process. Initially, we identified 3119 records, from which 7 duplicates were removed, 2977 were deemed irrelevant during the initial screening process, and 52 studies were excluded following full-text review. Finally, 83 studies employing MEPs for electrophysiological assessment were included in this review.

**Figure 1.**
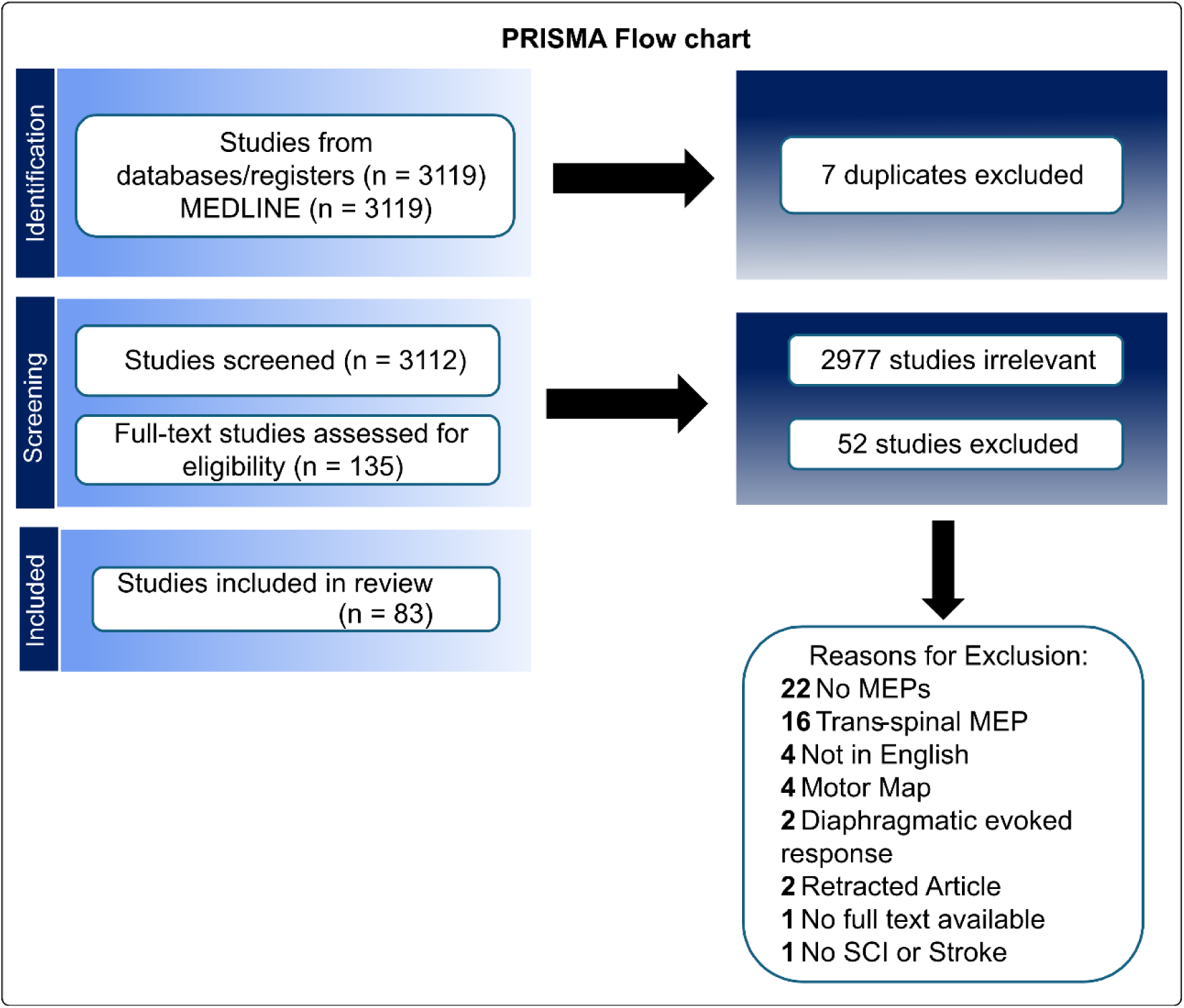
**PRISMA Flowchart.**

### 3.2 Characteristics of Reported Studies

**Table 1** summarizes the characteristics of studies included in this review, organized by post-injury intervention in rodent models of stroke and SCI. Studies were classified according to the type of post-injury intervention applied. Biomaterial-based interventions used engineered scaffolds or microenvironments to provide structural and biochemical support for tissue repair, often alongside cell delivery strategies. Cell and stem cell therapies were applied without additional scaffolds and aimed to promote motor recovery through cellular integration or paracrine effects. Drug and ligand treatments were differentiated based on whether the compounds were administered exogenously or produced endogenously by the animal, while combinatorial therapies included interventions combining multiple categories, reflecting a growing interest in multi-modal approaches. Neural stimulation interventions directly targeted central or peripheral nerves, including protocols such as direct cortical or peripheral stimulation, whereas studies employing intermittent theta-burst stimulation (iTBS) or transcranial magnetic stimulation (TMS) were grouped under TMS. Finally, studies with no post-injury intervention, focusing on endogenous plasticity, were categorized as not available (NA; **Figure 2a**).

**Figure 2.**
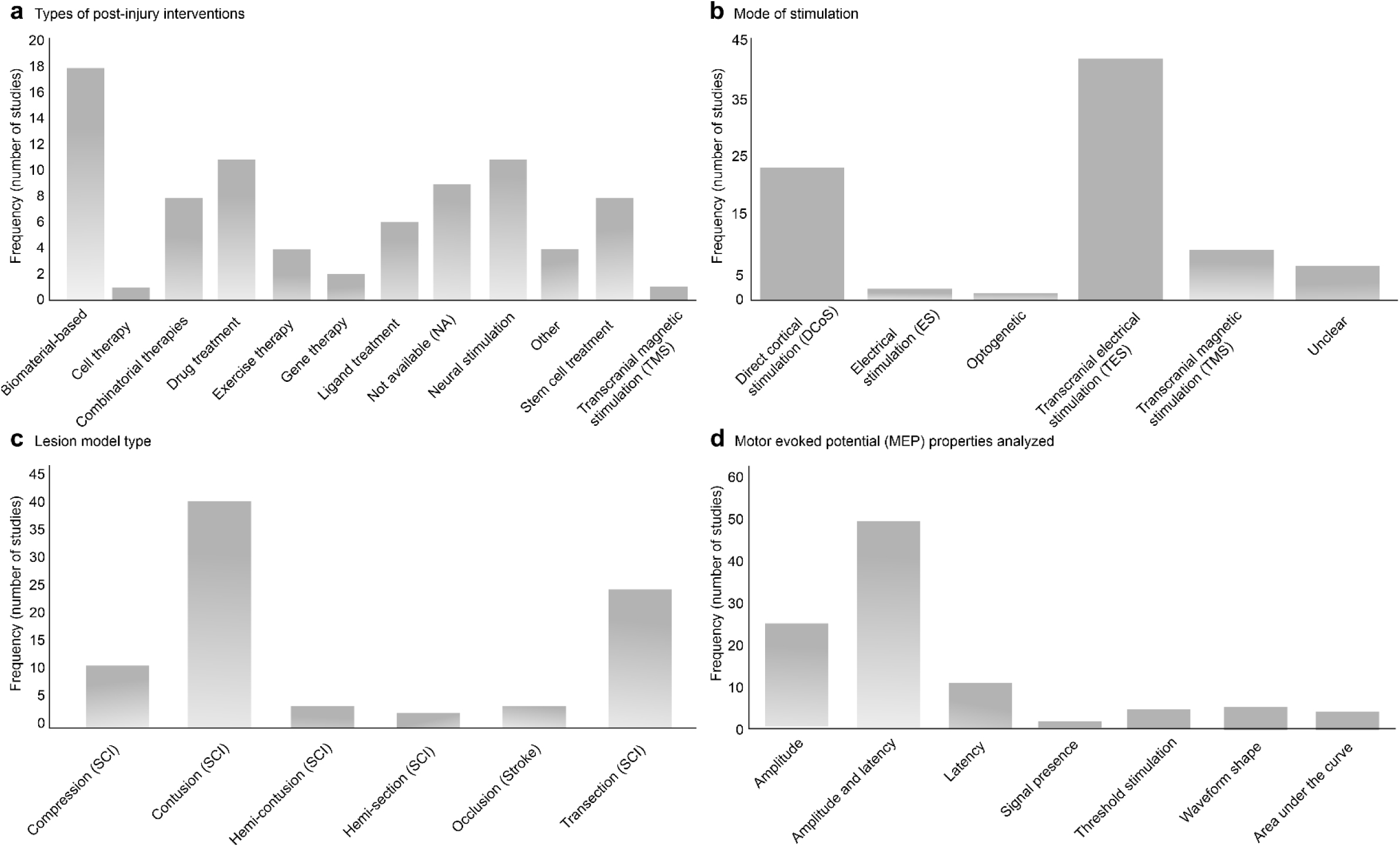
Characteristics of included studies. (**a**) Most MEP studies assess treatment effects; prominent interventions include biomaterials, pharmacologic agents, neural stimulation, and cell- or stem cell–based therapies. (**b**) Electrical stimulation is most commonly used to elicit MEPs. (**c**) SCI models predominate, with relatively few studies in stroke. (**d**) Amplitude and latency are the most frequently reported MEP metrics. *Note: NA (not available); NDP (No data provided); TMS (Transcranial magnetic stimulation); DCoS (Direct cortical stimulation); TES (Transcranial electrical stimulation); MEP (motor evoked potential); SCI (Spinal cord injury)*.

**Table 1.**
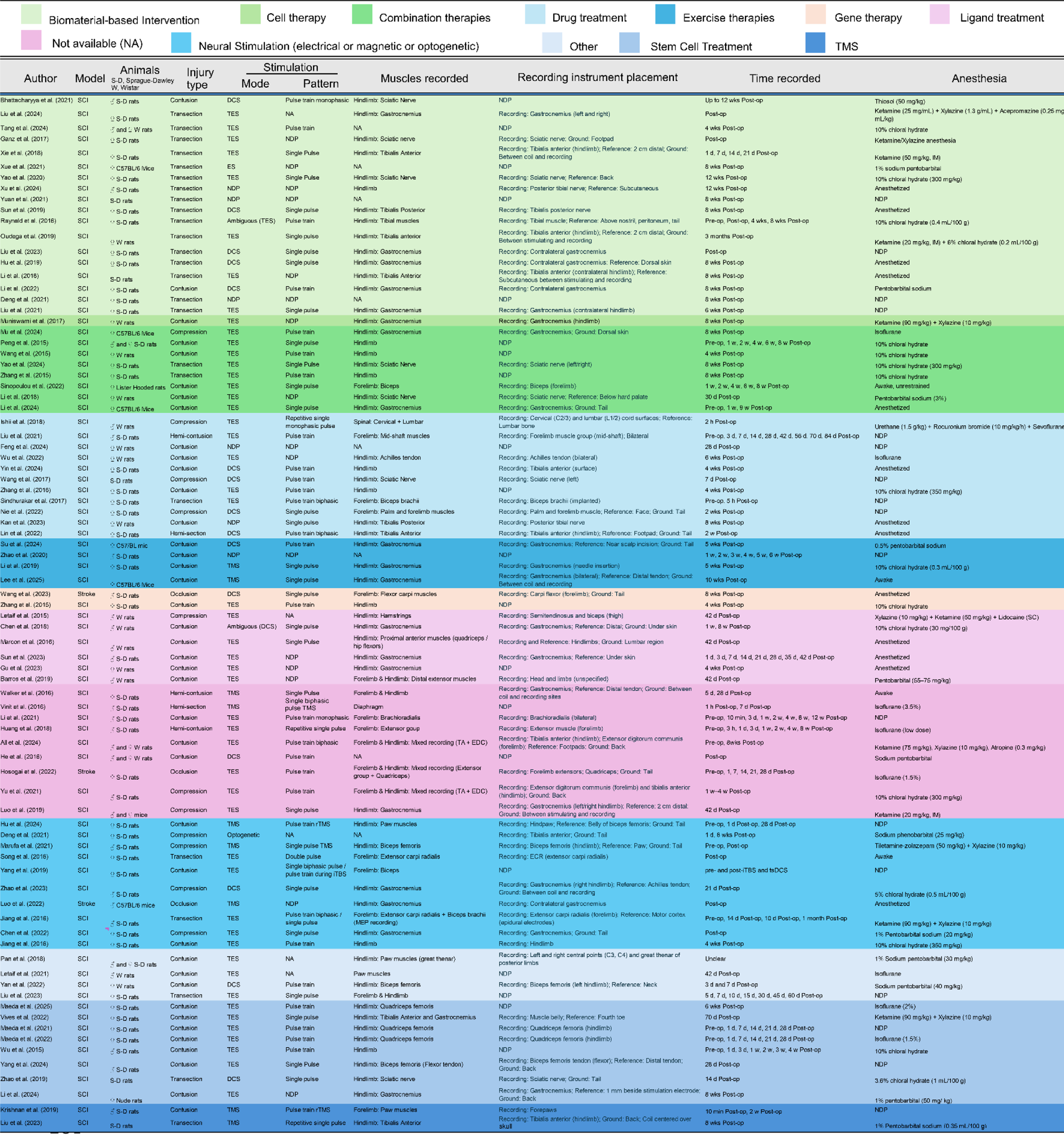
Summary of the characteristics of studies included in this review, organized by post-injury intervention in rodent models of stroke and SCI.

Across all 83 included studies, SCI models predominated, with only three employing experimentally induced stroke. Hindlimb muscles were the most frequently recorded, and stimulation protocols varied widely, including single-pulse, repetitive, and theta-burst patterns. Anesthetic approaches were similarly heterogeneous, representing a potentially large source of variability in MEP acquisition. These trends underscore the current emphasis on SCI models, the methodological diversity in stimulation and recording parameters, and the need for further stroke-focused research to validate MEPs as biomarkers of corticospinal tract integrity.

### 3.3. Modes of Cortical Stimulation Used Across Studies

Among the 83 publications reviewed, six cortical stimulation modalities were reported. Their usage frequency is illustrated in Figure 2b. Transcranial electrical stimulation (TES) was the most frequently used (42 studies, 50.6%), followed by direct cortical stimulation (DCoS; 23 studies, 27.7%) and transcranial magnetic stimulation (TMS; 9 studies, 10.8%). Optogenetic stimulation appeared rarely (1 study, 1.2%). Six studies (7.23%) provided ambiguous descriptions that prevented definitive identification of stimulation type and were classified as Unclear. Two studies (2.4%) employed electrical stimulation but were unclear as to whether it was transcranial or direct cortical stimulation and were categorized as electrical stimulation (ES).

### 3.4. Distribution of Stimulation Modes Across Lesion Types

Lesion type is an important consideration when designing MEP stimulation and recording protocols, as the specific injury can influence the interpretation of MEP properties such as amplitude and latency (19). Differences in lesion severity and location affect corticospinal tract integrity and can alter local electrical conduction, impacting signal transmission and recorded responses.

As seen in **Figure 2c**, lesion types were distributed as follows: contusion (n = 40, 48.2%), transection (n = 24, 28.9%), compression (n = 11, 13.3%), hemi-contusion (n = 3, 3.6%), hemisection (n = 2, 2.4%), and vascular occlusion (n = 3, 3.6%).

### 3.5. MEP Properties

**Table 2 and Figure 2D** summarize the motor evoked potential (MEP) properties reported across studies, organized by stimulation type. Categories are not exclusive; studies reporting multiple measures are listed next to each in the table. Across all modalities, amplitude and amplitude combined with latency were the most frequently measured MEP parameters reported in 25 and 49 studies, respectively. In contrast, other parameters such as Latency (11 studies), Area Under Curve (4), Threshold Stimulation (4), Waveform Shape (4), and Signal Presence (1) were reported far less frequently. Overall, this trend indicates that MEP amplitude, alone or in combination with latency, was the predominant outcome used to assess corticospinal excitability and motor recovery across experimental paradigms.

**Table 2.**
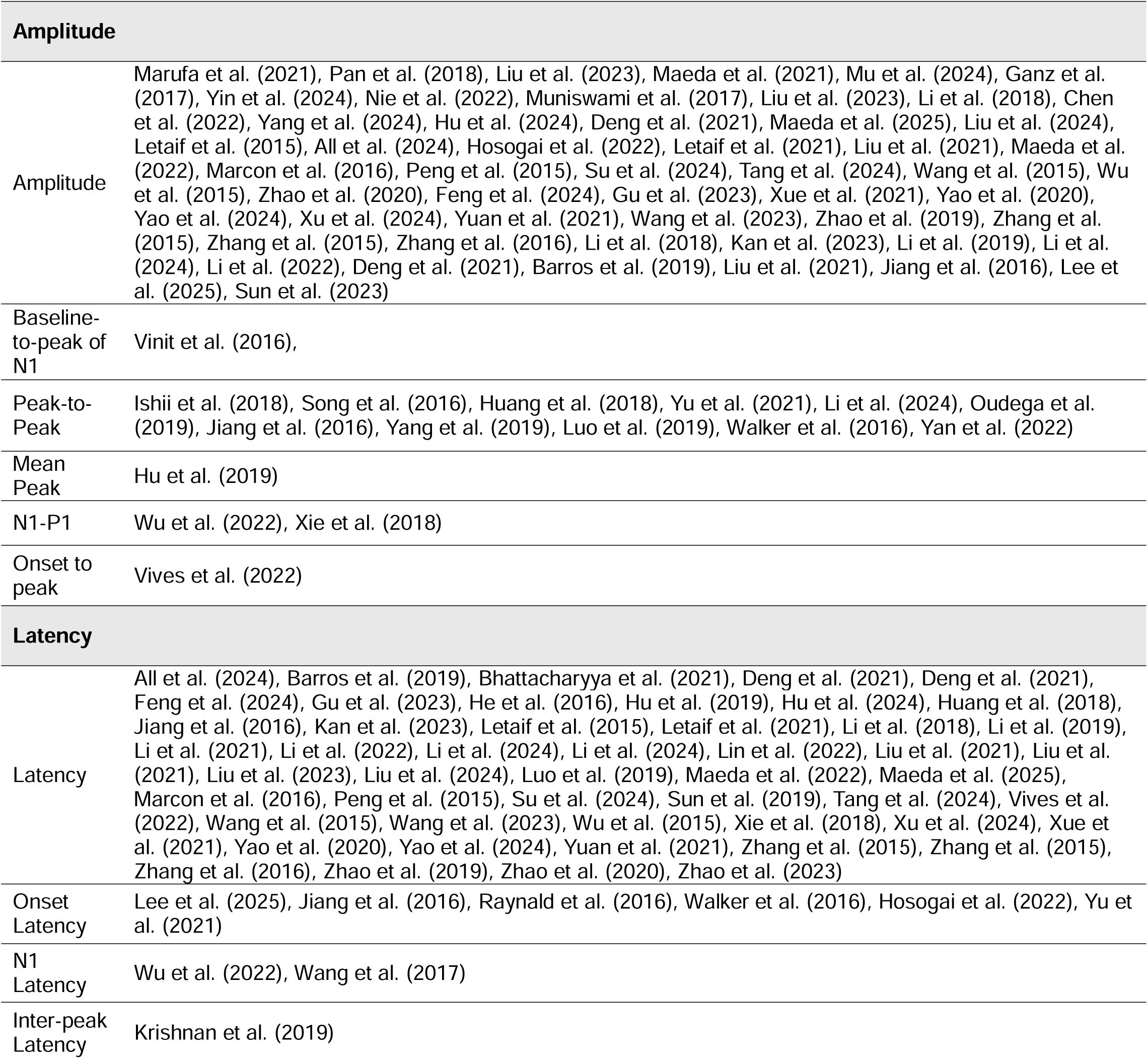
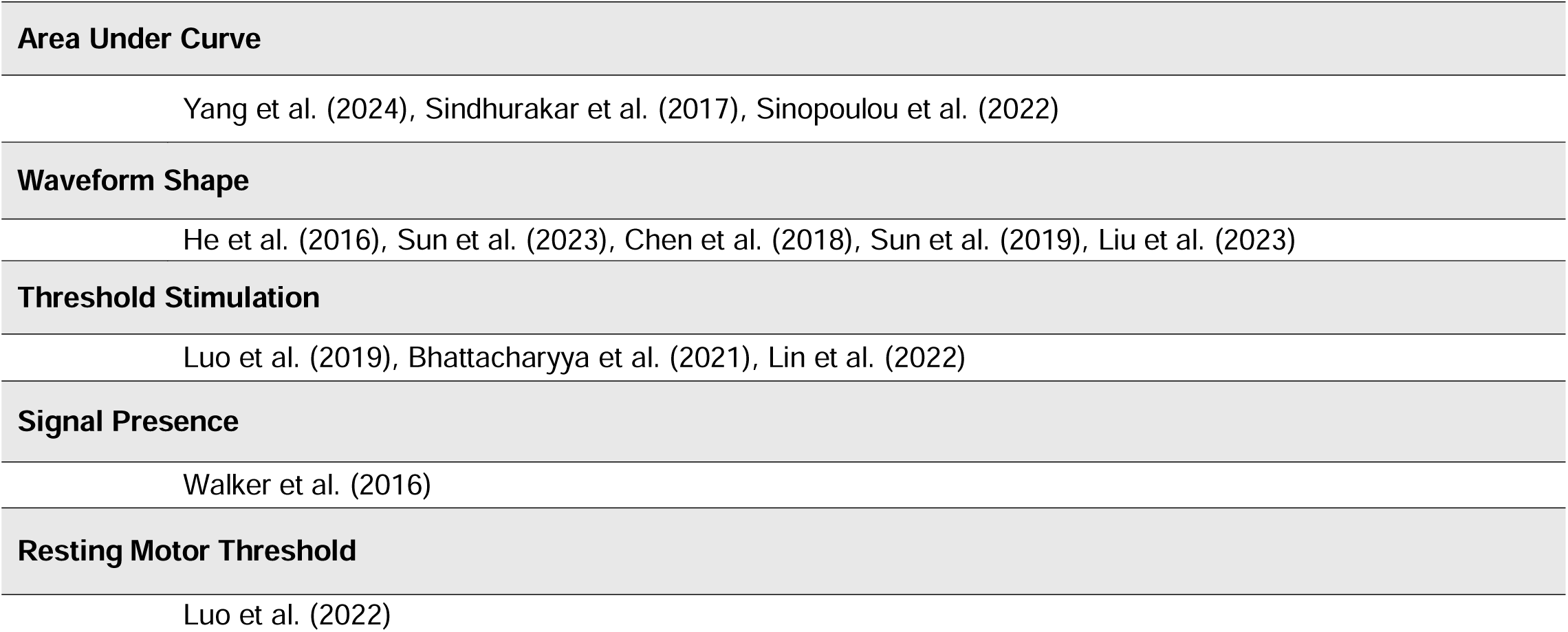
Summary of the motor evoked potential (MEP) properties reported across studies.

| Amplitude |  |
| --- | --- |
| Amplitude | Marufa et al. (2021), Pan et al. (2018), Liu et al. (2023), Maeda et al. (2021), Mu et al. (2024), Ganz et al. (2017), Yin et al. (2024), Nie et al. (2022), Muniswami et al. (2017), Liu et al. (2023), Li et al. (2018), Chen et al. (2022), Yang et al. (2024), Hu et al. (2024), Deng et al. (2021), Maeda et al. (2025), Liu et al. (2024), Letaif et al. (2015), All et al. (2024), Hosogai et al. (2022), Letaif et al. (2021), Liu et al. (2021), Maeda et al. (2022), Marcon et al. (2016), Peng et al. (2015), Su et al. (2024), Tang et al. (2024), Wang et al. (2015), Wu et al. (2015), Zhao et al. (2020), Feng et al. (2024), Gu et al. (2023), Xue et al. (2021), Yao et al. (2020), Yao et al. (2024), Xu et al. (2024), Yuan et al. (2021), Wang et al. (2023), Zhao et al. (2019), Zhang et al. (2015), Zhang et al. (2015), Zhang et al. (2016), Li et al. (2018), Kan et al. (2023), Li et al. (2019), Li et al. (2024), Li et al. (2022), Deng et al. (2021), Barros et al. (2019), Liu et al. (2021), Jiang et al. (2016), Lee et al. (2025), Sun et al. (2023) |
| Baseline-to-peak of N1 | Vinit et al. (2016), |
| Peak-to-Peak | Ishii et al. (2018), Song et al. (2016), Huang et al. (2018), Yu et al. (2021), Li et al. (2024), Oudega et al. (2019), Jiang et al. (2016), Yang et al. (2019), Luo et al. (2019), Walker et al. (2016), Yan et al. (2022) |
| Mean Peak | Hu et al. (2019) |
| N1-P1 | Wu et al. (2022), Xie et al. (2018) |
| Onset to peak | Vives et al. (2022) |
| Latency |  |
| Latency | All et al. (2024), Barros et al. (2019), Bhattacharyya et al. (2021), Deng et al. (2021), Deng et al. (2021), Feng et al. (2024), Gu et al. (2023), He et al. (2016), Hu et al. (2019), Hu et al. (2024), Huang et al. (2018), Jiang et al. (2016), Kan et al. (2023), Letaif et al. (2015), Letaif et al. (2021), Li et al. (2018), Li et al. (2019), Li et al. (2021), Li et al. (2022), Li et al. (2024), Li et al. (2024), Lin et al. (2022), Liu et al. (2021), Liu et al. (2021), Liu et al. (2023), Liu et al. (2024), Luo et al. (2019), Maeda et al. (2022), Maeda et al. (2025), Marcon et al. (2016), Peng et al. (2015), Su et al. (2024), Sun et al. (2019), Tang et al. (2024), Vives et al. (2022), Wang et al. (2015), Wang et al. (2023), Wu et al. (2015), Xie et al. (2018), Xu et al. (2024), Xue et al. (2021), Yao et al. (2020), Yao et al. (2024), Yuan et al. (2021), Zhang et al. (2015), Zhang et al. (2015), Zhang et al. (2016), Zhao et al. (2019), Zhao et al. (2020), Zhao et al. (2023) |
| Onset Latency | Lee et al. (2025), Jiang et al. (2016), Raynald et al. (2016), Walker et al. (2016), Hosogai et al. (2022), Yu et al. (2021) |
| N1 Latency | Wu et al. (2022), Wang et al. (2017) |
| Inter-peak Latency | Krishnan et al. (2019) |

|  |
| --- |
| **Area Under Curve** |
| Yang et al. (2024), Sindhurakar et al. (2017), Sinopoulou et al. (2022) |
| **Waveform Shape** |
| He et al. (2016), Sun et al. (2023), Chen et al. (2018), Sun et al. (2019), Liu et al. (2023) |
| **Threshold Stimulation** |
| Luo et al. (2019), Bhattacharyya et al. (2021), Lin et al. (2022) |
| **Signal Presence** |
| Walker et al. (2016) |
| **Resting Motor Threshold** |
| Luo et al. (2022) |

The preference for Amplitude and Latency was particularly apparent in transcranial electrical stimulation (TES) and direct cortical stimulation (DCoS) studies.

TES studies demonstrated the greatest methodological diversity, with various amplitude definitions (e.g., peak-to-peak, baseline-to-peak, inter-peak) and additional parameters such as area under curve and response duration. Optogenetic stimulation and studies utilizing TMS predominantly quantified amplitude and latency, with relatively few incorporating waveform shape or threshold metrics.

Overall, the distribution highlights a clear preference for amplitude-based MEP quantification, underscoring limited analysis of secondary electrophysiological features such as waveform morphology and temporal dynamics.

A large majority of these provide no detail on exactly how amplitude was measured. This heterogeneity in measurement definitions and reporting practices complicates cross-study comparisons and meta-analytic synthesis, suggesting a need for consistent reporting guidelines for MEP characterization in preclinical models.

This suggests a methodological emphasis on amplitude-based quantification of MEPs, with fewer studies incorporating temporal (latency) or waveform-based metrics. The limited exploration of electrophysiological features highlights that many research groups are not taking full advantage of more sophisticated analytical tools of the MEP.

### 3.6. Anesthetic Information

Anesthesia protocols were reported in 54 of the 83 studies, while 16 studies (19.3%) did not report whether they used anesthetics (NDP), and 13 studies (15.7%) did not report the name/dosage of the anesthetic used in their experiment (**Figure 3**). Among general anesthetics, chloral hydrate was most frequently applied (n = 16, 19.3%), followed by pentobarbital in its various forms (n = 11, 13.3%) and isoflurane (n = 8, 9.6%). Sodium phenobarbital was reported in one study (1.2%). Ketamine alone was used in two studies (2.4%), whereas ketamine-based combinations—including ketamine paired with xylazine, atropine, acepromazine, chloral hydrate, or lidocaine—were employed in nine studies (10.8%). Less commonly reported general anesthetics included urethane (n = 1, 1.2%), tiletamine-zolazepam (n = 1, 1.2%), and thiosol (n = 1, 1.2%). MEP recordings were conducted in awake and restrained animals in three studies and awake and freely moving in one (4.8% total).

**Figure 3.**
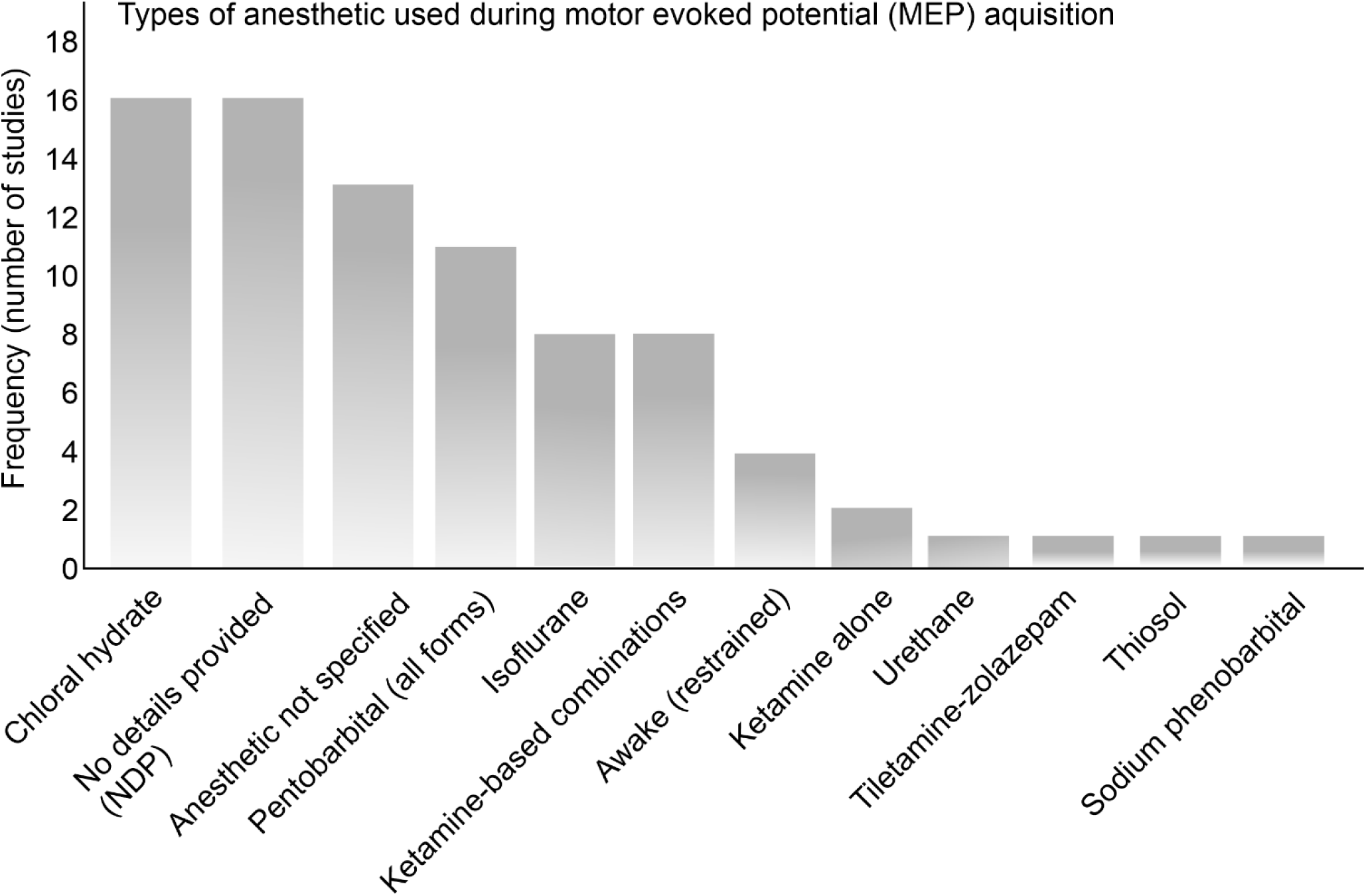
Types of anesthetics used during MEP acquisition.

### 3.7. Reporting Completeness and Methodological Transparency

To further evaluate the reproducibility of current MEP research practices, each study was systematically assessed for the completeness of methodological reporting. A scoring framework encompassing eight protocol components—ranging from stimulation and recording details to signal acquisition parameters—was applied to quantify reporting transparency across studies.

Scores ranged from 1 to 8, with a maximum of 8 representing full reporting across all methodological domains. The distribution was left-skewed, with most studies providing moderate-to-high detail (**Figure 4a**). The mean completeness score was 6.05 ± 1.25, indicating that, on average, two protocol components were omitted per study. A plurality of studies (44.6%) achieved scores of 7, while only 10 studies (12%) provided complete methodological descriptions for all eight categories. A small subset (8.4%) scored <4, reflecting low reporting transparency.

**Figure 4.**
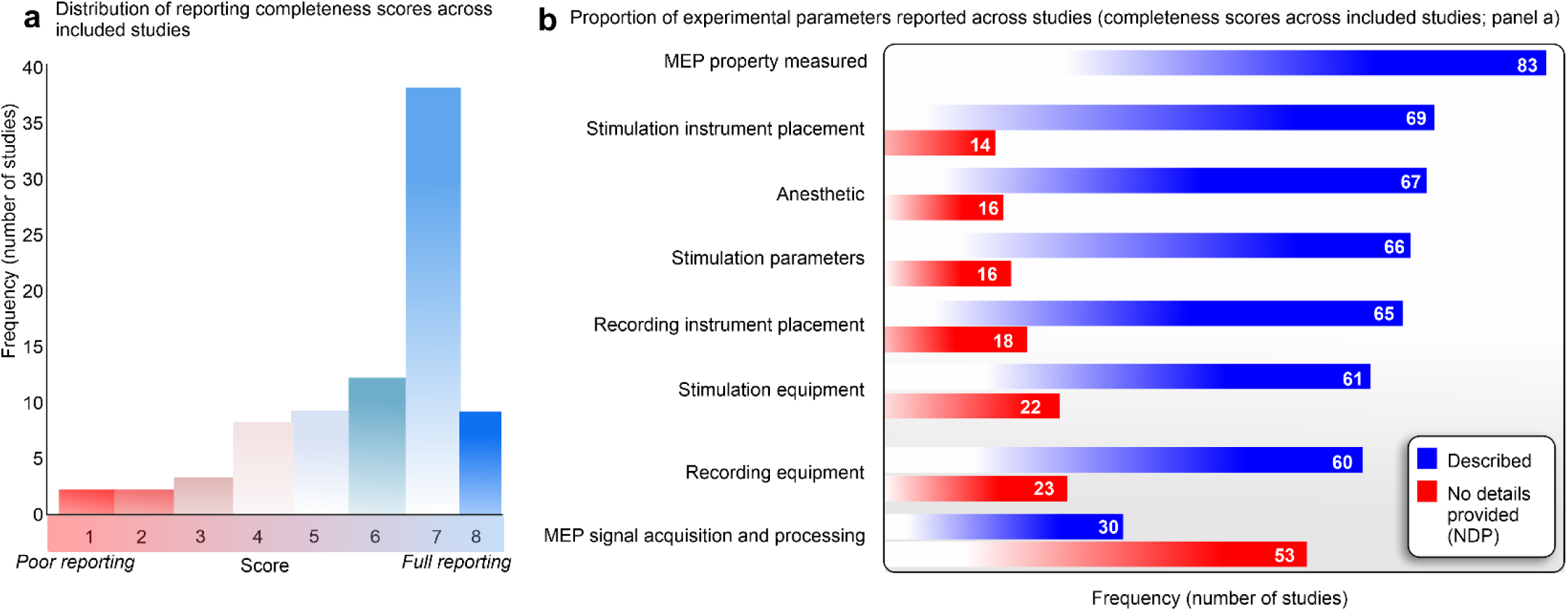
Reporting Completeness, Methodological Transparency, and Anesthetic Information. (**a**) Reporting Completeness and Methodological Transparency. (**b**) Detailed breakdown of items evaluated in the reporting completeness scores. *NDP (No data provided); MEP (motor evoked potential)*.

A detailed breakdown of reporting by parameter is presented in **Figure 4b**. The MEP property measured was reported universally in all 83 studies analyzed (100%), followed by stimulating instrument placement (83%), anesthetic information (82%), stimulation parameters (80%), recording instrument placement (78%), and equipment details for recording or stimulation (72–74%). In contrast, signal acquisition and processing parameters were the least frequently reported (36%).

### 3.8. Other study characteristics

Although CST projections are crucial for controlling hand and upper-extremity movements, most rodent studies have focused on the lower limbs. Of 83 studies, 11 (13.2%) examined forelimb/paw MEPs only, 54 (65.1%) investigated hindlimb MEPs, 6 (7.2%) recorded from both the forelimb and hindlimb, and 12 (14.5%) did not specify the recorded muscles.

A range of stimulation timing was used, including pulse trains and single-pulse paradigms.

## 4. Discussion (3730 words)

### 4.1. MEPs in Rodent Models of SCI and Stroke: A Preclinical Overview

The primary applications of MEPs observed across these studies can be broadly categorized as follows: treatment efficacy, lesion characterization and modelling, longitudinal monitoring and correlation with behavioural outcomes.

Numerous studies report positive effects of interventions on MEP parameters, typically manifesting as increased MEP amplitudes—signifying enhanced nerve conduction, greater recruitment of motor units, or stronger descending drive (20–29). Additionally, decreased MEP latency indicates faster conduction velocity along the motor pathways. Specific examples include rTMS leading to higher wave amplitude and lower latency (20), optogenetic stimulation promoting functional recovery and improving MEPs (21), and various cell therapies resulting in increased MEP amplitudes and/or decreased latencies (22,24,30).

#### 4.1.1. Experimental Applications of MEPs in Rodent Models of Stroke & SCI

As summarized in Table 1, MEPs are employed as electrophysiological tools to assess motor recovery and corticospinal integrity across a broad range of experimental studies. While MEPs are also employed in studies aiming to develop or improve existing models of spinal cord injury (11,31–36), they are primarily employed to assess the post-SCI intervention efficacy. Across these applications, MEPs provide a quantitative readout of descending motor pathway integrity that complements behavioural, histological, and molecular assessments.

The most prominent use of MEPs is to quantify the effects of various interventions aimed at promoting recovery after SCI or stroke. MEPs are used to gauge the impact of drug treatments on motor function (29,37–44), the efficacy of cell transplantation, including mesenchymal stem cells, oligodendrocyte precursor cells, and neural stem cells (22,24,30,45–49). Similarly, studies frequently use MEPs to evaluate nerve regeneration and functional improvement following the implantation of biomaterials or scaffolds (50–56).

MEPs are critical for evaluating the effects of electrical neuromodulation (13), repetitive transcranial magnetic stimulation (rTMS), trans-spinal magnetic stimulation, optogenetic stimulation, and other neuromodulation approaches (21,26,45,57–64). The benefits of interventions such as walking training or scar tissue resection are also monitored via MEPs (65–67).

MEPs occupy an intermediate position between cellular-level repair mechanisms and overt behavioural recovery, providing a functional readout of descending motor pathway integrity.

### Cell- and Stem Cell-Based Interventions

Stem cell research remains a major focus in experimental stroke and SCI studies, driven by the potential for cell-based therapies to promote repair through plasticity, remyelination, and regeneration. However, translation from rodent models to clinical application has proven difficult, emphasizing the need for mechanistic tools capable of linking cellular interventions to functional circuit-level outcomes. In this context, MEPs are frequently incorporated to assess whether stem cell–mediated changes result in restored CST conduction.

Across the literature reviewed, multiple studies explicitly employed MEPs to characterize electrophysiological changes following stem cell transplantation. For example, Maeda et al. (2021) examined longitudinal MEP changes in a rat SCI model following transplantation of mesenchymal stem cells (MSCs) derived from different tissue sources, using MEP amplitude and latency to compare recovery trajectories across treatment groups (68). Wu et al. (2015) investigated erythropoietin gene– modified neural stem cells delivered into the subarachnoid cavity, incorporating MEPs to probe protective effects on descending motor pathways (69). More recently, Yang et al. (2024) used MEP recordings to assess recovery following administration of hypoxic-preconditioned MSC-derived extracellular vesicles, linking changes in motor pathway conduction to astrocytic phenotype modulation via the miR-21/JAK2/STAT3 pathway (70).

Zhang et al. (2015) investigated whether edaravone combined with Schwann cell transplantation improved motor recovery post-SCI more than Schwann cell transplantation alone, finding that MEP latencies significantly shortened and amplitudes significantly increased in both treatment groups, with superior results in the combination group (71).

Stem cell therapies are also frequently embedded within combinatorial intervention designs. MEPs are used in studies pairing stem cells with pharmacological agents, neural stimulation, or biomaterial scaffolds to determine whether combined approaches yield additive or synergistic effects on CST function. For instance, stem cells loaded into electroconductive hydrogels or combined with cortical or spinal stimulation paradigms are evaluated using MEPs to assess functional connectivity and conduction across injured segments (51,53,56,72). These applications illustrate how MEPs enable researchers to dissociate cellular survival or integration from meaningful restoration of motor pathway function.

### Gene Therapy–Based Interventions

Gene therapy represents a more targeted experimental strategy aimed at modulating molecular pathways underlying synaptic plasticity and circuit reorganization. Although relatively few studies incorporate MEPs in gene therapy paradigms, their use highlights the sensitivity of electrophysiological measures for detecting pathway-specific functional changes.

Nie et al. 2022 used MEPs to explore the role of CDK-1 in chronic SCI and the effectiveness of CDK1 shRNA lentivirus injection to promote motor recovery, observing that MEP amplitude was increased gradually in the CDK1 shRNA group, approaching a normal MEP signal curve without full restoration (73).

Wang et al. (2023) provided a representative example by examining the effects of combined intramuscular AAV-BDNF and intranasal AAV-TrkB administration in a rodent stroke model. MEPs were used to assess central motor pathway conduction, serving as a functional bridge between enhanced neurotrophic signalling, synaptic plasticity of corticospinal connections, and behavioural recovery (74). In such studies, MEPs play a critical role in determining whether gene-driven molecular changes translate into improved systems-level motor output.

### Ligand and Drug–Based Interventions

Pharmacological and ligand-based treatments constitute one of the most common experimental applications of MEPs in rodent models of stroke and SCI. These studies use MEPs to evaluate neuroprotection, excitability modulation, conduction recovery, and optimization of treatment parameters across a diverse range of compounds.

Several investigations leverage MEPs to assess endogenous or physiologically relevant ligands. Letaif et al. (2015) examined the effects of estrogen following SCI, reporting a 17-fold decrease in MEP latency and a 7-fold increase in amplitude in treated animals, indicative of enhanced conduction velocity and excitability (35). Similarly, Chen et al. (2018) investigated bone morphogenetic protein 7 (BMP7), documenting not only changes in amplitude and latency but also alterations in MEP waveform morphology from a classic P1-N1-P2 to an M-shaped bimodal wave over time (75). The emergence of bimodal or M-shaped waveforms during recovery highlights a unique advantage of electrophysiological assessment: the ability to capture qualitative features of signal transmission that may reflect circuit reorganization and plasticity beyond what behavioural measures can detect.

Sindhurakar et al. (2017) used MEPs to determine if 4-Aminopyridine at clinically relevant levels alters the excitability of intact motor circuits, and found that cortical excitability and MEP enhancements remained significantly stronger beyond 2 hours after infusion (76).

MEPs are also widely used to optimize drug dosing and administration timing. Ishii et al. (2018) employed MEP recordings to compare edaravone dosing regimens and treatment windows following SCI, demonstrating that moderate dosing (3 mg/kg/h) and early administration produced superior electrophysiological recovery, whereas recovery outcomes diminished with higher doses (10 mg/kg/h) (77). In this context, MEPs functioned not only as efficacy markers but also as tools for refining experimental and therapeutic design.

More recent work extends pharmacological evaluation to compounds derived from traditional or plant-based medicine. Yin et al. (2024) used MEP amplitude to assess the effects of ginsenoside Rg1 on angiogenesis and functional recovery after SCI. They reported a markedly elevated amplitude in the rat group treated with ginsenoside Rg1, illustrating the applicability of electrophysiological assessment across diverse therapeutic traditions and mechanistic targets (27).

### Biomaterial-Based Interventions

Biomaterial-based approaches represent the most common category of MEP application in the literature reviewed and are particularly prevalent in transection injury models. These interventions aim to modify the lesion microenvironment, provide structural guidance for regenerating axons, and promote reconnection across disrupted pathways. In such models, the re-emergence of MEPs is often interpreted as evidence of restored functional continuity across the injury site.

Oudega et al., 2019 used MEPs to determine if a tube holding chitosan carriers loaded with neurotrophin-3 could trigger de novo neural tissue generation and functional recovery after insertion into a transection gap in the adult rat spinal cord. While MEPs were not detected in untreated SCI rats, they were in treated SCI rats, with smaller amplitudes but similar latencies to MEPs in intact controls (78).

Sun et al. 2019 used MEPs to estimate the therapeutic effects of an engineered collagen-chitosan scaffold fabricated by 3D printing technology on motor function recovery and nerve fiber regeneration. They observed decreased MEP latency and increased MEP amplitude in the treatment group compared to control groups (79).

Liu et al. (2023) evaluated an injectable, self-healing, electroconductive hydrogel loaded with neural stem cells and donepezil, using MEP amplitude to compare recovery across treatment combinations (80). Graded increases in amplitude across groups demonstrated how material properties, pharmacological support, and cell-based therapies interact to enhance nerve conduction. Xie et al. (2018) employed sodium hyaluronate–CNTF scaffolds to bridge gaps in spinal cord continuity. While the control group evoked no MEPs, the sodium hyaluronate-CNTF scaffold ground showed recovery after week 4, and demonstrated delayed but partial restoration of MEPs weeks after SCI (55). Critically, re-transection of the regenerated spinal cord abolished MEPs, providing strong electrophysiological evidence that recovered signals depended on newly formed descending connections rather than peripheral compensation alone.

Ganz et al. (2017) aimed to investigate the efficacy of an implanted 3D construct with oral mucosa-derived cells for functional recovery post-complete transection, finding that propagated MEPs in the induced group were significantly higher than the acellular group, suggesting partial connectivity, and MEP propagation was completely abolished after re-transection, indicating travel through CST across the lesion (81).

Other biomaterial studies incorporate MEPs to assess attenuation of secondary injury processes. Bhattacharyya et al. (2021) combined an iron oxide nanoparticle– impregnated hydrogel with magnetic field exposure to modulate the post-injury microenvironment, using MEPs as an integrative measure of cumulative effects on motor pathway function (82).

### Neural Stimulation–Based Interventions

MEPs are frequently used in studies involving neural stimulation, including electrical, magnetic, and optogenetic approaches. These paradigms directly target excitability and activity-dependent plasticity within motor circuits, making electrophysiological assessment particularly relevant.

Liu et al. 2023 used MEPS to explore the efficacy of a new non-invasive intermittent theta burst brain stimulation protocol to promote axonal regeneration and motor function repair following severe spinal cord transection; the study found that the treatment significantly increased the response amplitude of MEPs (83).

Yang et al. (2019) used MEPs as a physiological biomarker to confirm neurophysiological changes by theta burst brain stimulation and DCoS, observing a significant increase in peak-to-peak EMG amplitude of MEPs after both treatments, indicating enhanced corticospinal transmission (84).

Zhao et al. (2023) investigated double-target neural circuit-magnetic stimulation for promoting motor function recovery post-SCI, observing that while MEP waveforms were poor and latencies longer in the SCI, the treatment significantly shortened latencies, suggesting recovery of the descending motor pathway (85).

Deng et al. (2021) demonstrated that optogenetic activation of glutamatergic neurons in the primary motor cortex enhanced MEP amplitudes and shortened response latencies 6 weeks after SCI, suggesting strengthened descending conduction (21). By contrast, control stimulation paradigms failed to produce similar electrophysiological changes, highlighting the specificity of neurotransmitter-defined circuit activation. Jiang et al.(2016) used MEPs to investigate interactions between proprioceptive afferents and corticospinal axons, illustrating how changes in sensory input can reshape motor output through competitive circuit remodelling (86). These studies emphasize that recovery is not simply the restoration of pre-injury pathways, but the rebalancing of interacting motor and sensory circuits.

### Exercise, Physical Interventions, and others

MEPs are also applied in rehabilitation-focused studies. Zhao et al. (2020) used transcranial electrical MEPs to evaluate the effects of treadmill training onset after SCI, demonstrating that early training accelerated electrophysiological and behavioural recovery (65). They found that spontaneous recovery plateaued at the end of the 4^th^ week, and that treadmill training induced a rapid acceleration of related recovery in the early phase of recovery. Such findings suggest that rehabilitative activity and training timing can shape motor circuit plasticity and electrophysiological recovery after SCI.

Other studies employ MEPs to assess physical interventions, such as scar tissue resection, or to validate injury model refinements. For example, Liu et al. (2023) demonstrated changes in waveform amplitude and latency following scar resection at different time points, using electrophysiological outcomes to identify optimal intervention windows (67). Electroacupuncture-based combinatorial therapies further illustrate how MEPs are used to assess integrated treatment strategies that combine pharmacological, physical, and neuromodulatory elements (87).

Sinopoulou et al. (2022) used MEPs to evaluate temporal changes with a combinational treatment involving cortical epidural stimulation, rehabilitation and chondroitinase ABC, and found increased MEPs in the triple treatment group, supporting recovery of upper limb function (88).

Pan et al. 2018 used MEPs to evaluate post-SCI nerve conduction function if preceded by hyperbaric oxygen preconditioning, finding that MEP amplitude progressively decreased in SCI (89).

#### 4.1.2. Lesion Characterization and Modelling

Many studies employ MEPs to characterize the onset, progression, and severity of injury models. This is essential for validating preclinical models and understanding the natural history of recovery (11,36,90). MEPs help confirm lesion-induced deficits and track spontaneous recovery over time. A universal finding across SCI models is a significant reduction or complete absence of MEPs immediately post-injury (11,56,77,91,92). This initial suppression confirms the disruption of motor pathways. The type and severity of SCI or stroke influence MEP expression. Graded contusion models, for example, demonstrate severity-dependent MEP reductions (36). Unilateral lesions often lead to differential effects on ipsilateral versus contralateral limbs (32,33). MEPs have also been shown to be sensitive to ischemic changes (93).

Li et al. 2021 used MEPs to determine the correlation between electrophysiological testing and gross and motor function recovery following mild or severe SCI, revealing significant differences between mild and severe SCI conditions, and a progressive recovery of MEP activity from complete elimination post-SCI, with better recovery in mild cases (94).

Wang et al. (2017) addressed the involvement of autophagic activity in motor function recovery post-SCI, finding that MEP N1 wave peak latency was significantly longer with autophagy inhibitor 3-Methyladenine but significantly shorter with rapamycin treatment, suggesting autophagy’s role in attenuating SCI-induced spastic paralysis (95).

Letaif et al. (2021) used MEPs to standardize a model of intradural injection in experimental SCI in rats, with findings showing variable lower limb MEP latency and amplitude across groups, and experimental SCI groups having worse MEP results, though dural puncture or intrathecal injection did not worsen isolated SCI effects (96).

MEPs provide a valuable means for non-invasive (or minimally invasive) longitudinal assessment of motor pathway integrity, enabling researchers to track changes over time in response to injury or intervention (32,33,68,91,97–99). Many studies seek to correlate MEP findings with behavioural functional recovery, using metrics like the Basso Mouse

Scale or other motor tests (28,29,32,33,35,37,38,41,47,51,61,62,65–67,70,90,100–103). Subsequent recovery is often characterized by the reappearance of MEPs or increases in amplitude, which may indicate spontaneous repair or the success of therapeutic interventions. This helps bridge electrophysiological measures with tangible functional improvements.

While many studies demonstrate a positive correlation between improved MEPs and better behavioral scores, suggesting MEPs are a good proxy for functional recovery (32,66), some studies highlight a more nuanced relationship. For instance, All et al. (2024) (11) specifically noted that Basso Mouse Scale scores and MEP signals were not necessarily related in their model, suggesting that different physiological processes might underpin behavioral versus electrophysiological recovery in certain contexts.

#### 4.1.3. CST, upper extremity control and proportional recovery

In thoracic and lumbar SCI models, hindlimb MEPs provide a direct assessment of descending motor transmission through or below the injured spinal segments. By contrast, forelimb MEPs may provide a complementary measure of supralesional changes. For example, recording forelimb MEP (above the lesion) captures preserved corticospinal and brainstem-descending drive to muscles and thus provides a sensitive index of supraspinal excitability and plasticity after thoracic or lumbar spinal cord injury (104). Monitoring these responses distinguishes central (cortical or subcortical) adaptations from local spinal changes and therefore is essential for interpreting how brain motor networks respond to, and potentially compensate for, caudal lesions. Despite the CST’s central role in hand and upper-extremity function (1), most MEP studies in rodents have focused on the lower limbs. It is known from clinical studies that upper-extremity deficits commonly arise from disrupted CST signalling (MEP-positive post-stroke survivors) (4,105), and that residual CST integrity strongly predicts motor recovery (1,3,4). However, recent proportional recovery studies in rats (106,107) and mice (108) have not incorporated forelimb MEPs. Incorporating forelimb MEP recordings into rodent upper-extremity studies may therefore strengthen the use of CST integrity as a functional biomarker of motor recovery, particularly when the experimental focus is skilled forelimb control, supralesional plasticity, or CST-dependent recovery mechanisms.

### 4.2. Methodological Considerations for MEP Assessment in Preclinical SCI and Stroke

While MEPs are of great interest for assessing recovery after SCI and stroke, their application presents several methodological considerations. Although this electrophysiological assessment offers high sensitivity, it is also notably susceptible to artifacts at the brain stimulation and MEP recording sites and to the type of anesthetic used. These methodological sources of variability are closely linked to a second challenge: the interpretation of MEP characteristics themselves. Before considering stimulation, recording, and anesthesia protocols, it is therefore important to clarify how commonly reported MEP features have been interpreted physiologically across the included studies.

#### 4.2.1 MEP Characteristics and Their Physiological Interpretation

Motor evoked potentials have the potential to be utilized as functional biomarkers for corticospinal tract integrity. As noted earlier, they provide several advantages over traditional imaging techniques because the generation of an MEP depends on successful activation and conduction through descending motor pathways, spinal circuitry, peripheral nerves, and the recorded muscle response. As a result, MEPs can provide a functional readout of neurophysiological recovery of descending pathways and may help bridge underlying physiological changes with observable behavioural outcomes, making them useful for tracking plasticity and treatment efficacy in the nervous system.

This potential, however, depends on whether changes in MEP characteristics can be meaningfully linked to physiological changes. Difficulty in interpretation remains one of the greatest obstacles to their widespread use. Given this interpretive uncertainty, examining how MEP characteristics are interpreted across different experimental applications may help clarify how individual MEP properties have been represented in recent studies and highlight where methodological or interpretive improvements are needed.

### Amplitude

Amplitude was the most widely utilized MEP property for quantifying electrophysiological change across the studies analyzed in this review. It is commonly used as a readout of corticospinal excitability and descending motor output; however, it is also strongly shaped by stimulus intensity, anesthetic state, recording site, and signal quantification method, making it vulnerable to over-interpretation. The safest interpretation is therefore that amplitude represents the magnitude of the MEP under specific stimulation and recording conditions. It should be understood as a conditional marker of neural excitability and motor output, rather than as a standalone estimate of anatomical tract integrity.

Across the included studies, MEP amplitude was most often used as a functional marker of descending motor pathway conduction and recovery, particularly in spinal cord injury models. In transection studies, where the spinal cord is physically severed, the presence or increase of propagated MEP amplitude was generally interpreted as evidence of partial restoration of connectivity across the lesion site. For example, Ganz et al. (2017) used the presence of higher MEP amplitudes in cell-loaded scaffold animals, relative to acellular controls, as evidence of partial functional connectivity between the motor cortex and hindlimb motor neurons, although amplitudes remained below those observed in intact animals(81). The abolition of MEPs after rostral spinal cord re-transection further supported the interpretation that the recorded response depended on conduction across the lesion, rather than on spinal locomotor activity alone. Similarly, Liu et al. (2023) used increasing MEP amplitude over time as evidence of improved motor and nerve conduction recovery following a combined biomaterial, neural stem cell, and donepezil-loaded hydrogel intervention(80).

In compression SCI and non-SCI motor injury models, amplitude was also used to track electrophysiological recovery or plasticity, though its interpretation was less directly tied to structural reconnection. Hosogai et al. (2022), in a rodent stroke model, reported that MEP amplitude decreased acutely after injury and partially recovered over time, suggesting that amplitude can reflect spontaneous recovery-related plasticity even without an external intervention(97). Marufa et al. (2021) used amplitude to evaluate the effects of repeated theta-burst stimulation after spinal cord compression injury, reporting time-dependent amplitude changes across injury severities and later differences between stimulation protocols in the severe-injury group(58). Taken together, these studies suggest that MEP amplitude is most often interpreted as a functional recovery marker across rodent motor injury models, with particularly frequent use in SCI studies. However, because amplitude is influenced by stimulation intensity, anesthesia, recording site, and quantification method, it should be interpreted as a conditional electrophysiological marker rather than a direct measure of anatomical tract integrity.

### Latency

Latency was another frequently reported MEP property, used to quantify the timing of the evoked motor response rather than its magnitude. Broadly, MEP latency reflects the time required for the stimulation-induced response to reach the target muscle, including cortical activation, conduction through descending motor pathways, spinal motor neuron activation, and peripheral motor conduction. Because of this, latency is influenced by multiple levels of the motor system rather than by corticospinal tract conduction alone. It may also vary depending on stimulation parameters and the excitability state of the motor pathway; for example, voluntary contraction can shorten MEP latency by increasing spinal motoneuron excitability and lowering the threshold for response generation (109). Latency should therefore be interpreted as a composite measure of conduction timing and motor pathway excitability, rather than as a direct or isolated measure of anatomical tract integrity.

Across the included studies, prolonged MEP latency was generally interpreted as evidence of impaired conduction following injury, while reduced latency after intervention was treated as a marker of improved electrophysiological recovery. This pattern was observed across multiple SCI models and intervention types. In compression and ischemia/reperfusion injury models, longer latencies were reported after injury and were used as evidence of delayed motor pathway conduction. Interventions associated with shorter latencies, including drug treatment in Wang et al. (2017) and neural stimulation in Zhao et al. (2023), were interpreted as improving motor pathway function or promoting recovery of descending motor conduction(85,95). Similarly, in a transection model, Raynald et al. (2016) used shorter latency following a biomaterial-supported cell transplantation intervention as evidence of improved lower-extremity functional recovery(110).

However, latency was not always interpreted through a simple injury-worsens/intervention-improves framework. In a contusion SCI model, Krishnan et al. (2019) used latency to assess the effects of rTMS on corticospinal excitability, but stimulation was associated with longer latencies, particularly when applied acutely(111). This suggests that latency may capture stimulation-related changes in pathway recruitment or excitability that are not always straightforwardly equivalent to improved conduction. Taken together, these studies suggest that MEP latency is most often interpreted as a marker of conduction delay or recovery across rodent SCI models, including compression, contusion, and transection injuries. However, because latency reflects activity across cortical, spinal, and peripheral components of the motor pathway, and because studies vary in how latency is defined and measured, it should be interpreted as a composite electrophysiological marker rather than a standalone indicator of corticospinal tract repair.

### Waveform Shape

Waveform shape was a less commonly emphasized MEP property, but when reported, it was generally used to describe the overall morphology or recognizability of the evoked response. Unlike amplitude and latency, which quantify the size and timing of the MEP, waveform shape provides a more qualitative indication of how the response is organized. Changes in waveform morphology may reflect altered synchronization of descending volleys, changes in motor unit recruitment, disrupted conduction, or reorganization of the motor pathway after injury. However, waveform shape is also difficult to interpret in isolation, as MEPs can vary naturally across muscle groups and clinical populations. For example, MEPs recorded from non-hand muscles tend to be more polyphasic, and altered or more complex waveforms have also been reported in conditions such as amyotrophic lateral sclerosis, myoclonus dystonia, multiple sclerosis, and stroke (109). Therefore, waveform shape should be interpreted as a contextual marker of response organization rather than as a direct indicator of a single physiological mechanism.

Across the included studies, waveform shape was most often used to support interpretations of injury-related disruption or intervention-related recovery, particularly when paired with amplitude or latency. In contusion models, a typical MEP waveform was reported in sham or pre-injury conditions, while injury was associated with reduced, unclear, or altered waveforms. Ligand-based interventions, including BMP-7 treatment in Chen et al. (2018) and BMP-7-induced bone marrow mesenchymal stem cell treatment in Sun et al. (2023), were associated with later recovery of detectable MEP responses and changes in waveform morphology, alongside increased amplitude(75,112). In these studies, waveform shape was not interpreted as a standalone outcome, but rather as part of a broader electrophysiological pattern suggesting improved motor pathway function after intervention.

Waveform shape was also reported in transection models, where the reappearance or alteration of the evoked response was used to support evidence of functional recovery across a more severely disrupted pathway. Sun et al. (2019) included waveform morphology alongside amplitude and latency to evaluate a biomaterial-based scaffold intervention, interpreting the combined electrophysiological changes as evidence of improved neurological recovery(79). Similarly, Liu et al. (2023) reported detectable MEP waveforms after scar tissue resection, with accompanying changes in amplitude and latency over time(67). In these cases, waveform shape appeared to function mainly as evidence that an organized evoked response could be detected after severe injury or intervention, rather than as a precisely defined measure of anatomical repair.

Taken together, these studies suggest that MEP waveform shape is used as a supportive marker of electrophysiological organization and recovery, especially when injury disrupts the clarity or detectability of the evoked response. However, because waveform morphology can be influenced by recording muscle, injury model, motor pathway reorganization, and the degree of synchrony in the evoked response, its physiological meaning is less specific than amplitude or latency. It is therefore best interpreted alongside quantitative MEP properties, rather than as an independent measure of corticospinal tract integrity.

Although few included studies quantified waveform complexity in detail, features such as the number of peaks, temporal dispersion, and polyphasic structure may provide additional information about the synchrony of descending volleys, spinal motor neuron recruitment, and reorganization of motor circuitry after injury (109). Future rodent studies would benefit from reporting whether MEPs are mono- or polyphasic, whether peak timing changes after injury or intervention, and whether responses differ across recorded muscles or stimulation sites. These features may be particularly important when distinguishing cortically evoked responses from spinally or peripherally mediated responses. The use of more advanced machine learning techniques can help to quantify complex patterns in waveform shape in more detail.

### Area Under the Curve

Area under the curve was used less frequently than peak-to-peak amplitude, but it served a similar purpose as a measure of MEP size or overall evoked muscle activity. While amplitude captures the difference between the largest positive and negative peaks, area under the curve reflects the total magnitude of the rectified MEP response across a defined time window. This distinction is important because peak-to-peak amplitude is most valid when the waveform is relatively simple and does not contain polyphasic oscillations. When MEPs are more complex, area under the curve may provide a broader estimate of response size by capturing activity distributed across the waveform, rather than relying on a single peak-to-peak difference. Like amplitude, however, area under the curve remains dependent on stimulation intensity, recording conditions, signal processing, and the time window selected for analysis. It should therefore be interpreted as an integrated estimate of evoked muscle activity rather than as a direct measure of anatomical tract integrity.

Across the included studies, area under the curve was generally used to evaluate intervention-related recovery in SCI models. In contusion models, increased area under the curve was interpreted as evidence of improved electrophysiological function or greater evoked muscle activity after treatment. Yang et al. (2024), for example, reported higher MEP area under the curve and amplitude following hypoxic-preconditioned stem cell-derived extracellular vesicle treatment, interpreting these changes as improved hindlimb electrophysiological recovery(70). Similarly, Sinopoulou et al. (2022) used area under the curve and response duration to assess evoked muscle activity after a combinatorial intervention involving cortical stimulation, rehabilitation, and chondroitinase ABC (88). Increased evoked muscle activity in the treatment group was interpreted as supporting recovery of forelimb function after cervical contusion injury.

Area under the curve was also used in a transection-related injury model to assess drug-related changes in motor circuit excitability. Sindhurakar et al. (2017) used this measure to evaluate the effects of 4-aminopyridine and found enhanced physiological responses in injured animals, which was interpreted as strengthened spinal and cortical excitability after treatment (76). In this context, area under the curve was less directly framed as structural recovery and more as a marker of increased responsiveness within remaining or injured motor circuits.

Taken together, these studies suggest that area under the curve is used as an alternative or complementary measure of MEP size, especially when overall evoked activity may be more informative than a single peak-to-peak amplitude value. Its use may be particularly relevant when waveform morphology is complex or when studies aim to capture changes across stimulation intensities.

#### 4.2.2. Type of brain stimulation and electrode position

Placement was frequently over the skull, often aligned with anatomical landmarks like bregma and lambda to target the sensorimotor cortex (46,57,68,98). Specific coordinates were sometimes provided. Some studies simply mentioned placement over the motor cortex. Many studies specified electrode placement relative to bregma and lambda (38,57,68,74,92,113,114). Some provided specific distances (e.g., “AP + 0.00 mm, ML ± 2.00 mm, and DV 1.5 mm” (56), “2 mm posterior to the bregma, 2 mm lateral to the midline” (30), “AP: ±1.0 mm; ML: ±1.25 mm; bregma as the center” (68,113). Specific anatomical points like the anterior fontanel, coronal suture, and sagittal suture were often used as reference points for motor cortex stimulation (55,84,90,90,112). Many studies referred to the “motor cortex,” “sensorimotor cortex,” or “cerebral cortex” without providing explicit coordinates, relying on the general understanding of these regions (91,96,102,115,116). Descriptions like “over the vertex of the head” or “above the skull” were also common, possibly implying a broad area (60,95). Almost a fourth of the studies (22/83, 26.5%, Figure 4C) provided no details about stimulation electrode placement.

Several studies utilized specific TMS devices such as YIRUIDE (91), Magstim (57,62), and MagVenture (59,68,92,98). Coil types include specific animal coils (91), 5.5 cm doughnut-shaped coils (97), figure-of-eight coils (57,74,92), and circular coils (43,59,71).

Electrical stimulation involved various electrode types and placements. These included screw electrodes over the sensorimotor cortex (11,30,90,94,95,99,117–119), subdermal needle electrodes over the motor cortex (120), and electrodes on the skull surface (84,112,121). A methods study in mice (not in our sample) comparing some of these found less inter-session variability with scalp surface stimulation as compared to subcutaneous needles or epicranial screws (122). The same study also reported reduced variability with surface vs. intramuscular needle MEP recording.

Details on stimulation parameters varied widely, including intensity [e.g., 120% of resting motor threshold for TMS (59,91), single electrical pulse of 10mA (102)], frequency (e.g., 5 Hz, 10 Hz, 50 Hz for electrical stimulation), pulse width (e.g., 100 µs, 200 µs for electrical pulses), and number of pulses/trains (26,57,59,91,92,123). Some studies specified settings for continuous stimulation or single pulses.

In summary, while some studies offered precise coordinates or clearly defined anatomical landmarks, many others provided only general region descriptions or lacked this information entirely. Moreover, stimulation parameters varied widely. This variability in reporting and methodology could impact the reproducibility and direct comparison of findings across different studies.

#### 4.2.3. Electromyographic recording of the MEP

The description of EMG electrode placement varies substantially across the studies, from highly detailed to not provided at all.

Many studies precisely described the placement of EMG electrodes in specific muscles, often distinguishing between recording, reference, and ground electrodes. Common hindlimb sites included the *tibialis anterior* (29,30,37,38,54,68,84,121,123), *gastrocnemius* (85,112,118), and *biceps femoris* (36,45,60,69,70,100,124). For forelimb muscles, electrodes were mostly placed in the *extensor digitorum communis* (11,33) and *extensor carpi radialis* (47,100,125,117). Some studies directly recorded from peripheral nerves, such as the sciatic nerve (51,61,69,81,82,95,126). When provided, reference electrodes were typically placed on a paw, neck, distal tendon, hard palate, or subcutaneously. Ground electrodes were commonly placed on the tail or back of the rat, or subcutaneously between the stimulating and recording electrodes.

Other studies provided more general locations like “muscle groups in the mid-shaft of the forelimb” (59,65,96) or indicated that electrodes were “inserted intramuscularly” without naming specific muscles (33). About one in five did not offer any specific information regarding EMG electrode placement (18/83, 21.7%, Figure 4C).

In summary, there is a lack of information about precise placement of recording electrodes in many studies. Details about the exact location on the muscle and directions in relation to muscle fibres would increase reproducibility, similar to the guidelines used in human studies (e.g. SENIAN guidelines) (127). These guidelines help to ensure electrodes are in line with the orientation of the muscle fibres and away from the innervation zone. Placement of electrodes over innervation zones may cause dampened amplitudes and shifts in frequencies of the EMG (128).

The use of high-density EMG arrays can help identify innervation zones and provide a more comprehensive picture of the MEP distribution across muscle regions (129). This can also help in detecting changes in innervation zones due to neuroplastic changes during the recovery process after SCI and stroke (130).

#### 4.2.4. Anesthetics

The studies employed a variety of anesthetic agents and administration methods during MEP acquisition, with some experiments even conducted on awake animals. Surprisingly, isoflurane was used in 8 studies. It is known that isoflurane anesthesia severely hinders the acquisition of MEPs (131). Other anesthetics included pentobarbital sodium, chloral hydrate, ketamine and xylazine combinations. Some studies mentioned additional agents like atropine. Three studies explicitly stated that animals were awake and restrained. 13 studies provided no details for the anesthetic information (15.7%, **Figure 3**).

We recommend using Ketamine anesthesia, specifically a Ketamine/Xylazine/Atropine cocktail, as it yielded the most consistent and high-amplitude MEP signals in rats (131). In general, methodological studies outside of our sample have shown that different anesthetics have dose-dependent effects on MEP amplitude/latency (132,133) and interventions like deep brain stimulation (as measured by MEP) (134) and that effects on amplitude can sometimes vary systematically over time (135), arguing that where MEP amplitude/latency is important, dose and timing should be tightly controlled and reported in detail.

#### 4.2.5. Recruitment curves

Consistently, MEP amplitude (often peak-to-peak) and latency (onset to peak or first deflection) are the predominant quantitative measures across almost all studies. Other metrics like waveform shape, area under the curve, and threshold are also occasionally reported. Amplitude features extracted from MEPs can be influenced by various factors, including electrode positioning, anesthetic levels, and other experimental conditions. The use of comprehensive recruitment curves may help to mitigate this variability. Recruitment curves characterize the full input–output relationship of corticospinal excitability, capturing threshold, gain (slope), and maximal response, rather than a single-point amplitude that is highly placement- and state-dependent. In SCI, recruitment-curve metrics can reveal excitability changes missed by single-intensity measures; for example, a preliminary TMS study showed a steeper MEP recruitment curve in patients despite unchanged resting motor threshold, suggesting enhanced spinal/corticospinal responsiveness (136).

Methodological advantages include the mitigation of variability from electrode/coil placement, anesthesia, and day-to-day impedance by modelling responses across intensities (136,137). It also enables standardized, comparable metrics (e.g., S50— stimulation intensity that elicits 50% of the maximal evoked response on a recruitment curve, slope, MEP max, AUC-recruitment curve) and clearer reporting of stimulation intensity relative to motor threshold, which was often missing in the reviewed studies.

In many of the studies reviewed here, it was unclear at what stimulation intensity the MEPs were acquired (e.g., 120% of motor threshold), which represents a fundamental control. A recruitment curve, however, considers all stimulation intensities, from the motor threshold to the plateau of the maximal response. From this curve, several features can be extracted, such as the stimulation intensity required to achieve 50% of the maximum MEP response (S50) and the slope of the recruitment curve, among others.

Finally, the use of recruitment curves increases sensitivity to subtle treatment effects; recent work detected drug-related differences using recruitment-curve features that single-intensity amplitudes could miss. For example, we have recently used this approach to determine subtle differences with an AMPA agonist treatment (138).

Of the 83 reviewed studies, none described the use of features extracted from recruitment curves. We recommend that future studies follow best practices in utilizing recruitment curves to determine CST integrity and excitability.

## 5. Conclusion and future directions (225 words)

This systematic review highlights the indispensable role of MEPs in preclinical SCI and stroke research, offering a quantifiable window into motor pathway function. MEPs are crucial for characterizing injury models, monitoring disease progression longitudinally, and rigorously evaluating therapeutic interventions. By bridging the gap between cellular and molecular changes and observable behaviour, MEPs provide an integrated functional readout of neural circuitry. Despite their substantial value, the methodological landscape of MEP acquisition and interpretation in rodent models remains highly variable. This heterogeneity, particularly in stimulation protocols, anesthetic choices, and reporting of electrophysiological features, limits direct cross-study comparisons and hinders the translational impact of research findings. The consistent emphasis on amplitude and latency as primary metrics, while valuable, often overlooks the richer insights offered by more comprehensive analyses like recruitment curves.

### 5.1. Future directions

To enhance the robustness, reproducibility, and translational potential of MEP-based research in preclinical SCI and stroke studies, we strongly recommend the adoption of standardized protocols. Future studies should prioritize clear and detailed reporting of all methodological parameters, including specific stimulation intensities (e.g., as a percentage of motor threshold), precise electrode placements, and anesthetic dose and timing. Furthermore, we advocate for a more widespread integration of comprehensive recruitment curve analysis, as this approach captures a broader range of physiological information—such as the 50% maximal response and curve slope—which can reveal subtle changes not detectable by single-intensity measurements.

